# SimBit: A high performance, flexible and easy-to-use population genetic simulator

**DOI:** 10.1101/2020.05.12.086884

**Authors:** Remi Matthey-Doret

**Affiliations:** Department of Zoology and Biodiversity Research Centre, University of British Columbia, Vancouver, British Columbia V6T 1Z4, Canada; Institute of Ecology and Evolution, University of Bern, CH-3012 Bern, Switzerland

## Abstract

SimBit is a general purpose and high performance forward-in-time population genetics simulator. SimBit has been designed to be able to model a wide diversity of complex scenarios from a simple set of commands that are very flexible. SimBit also comes with a R wrapper that simplifies the management of an entire research project from the creation of a grid of parameters and corresponding inputs, running simulations and gathering outputs for analysis. Implementing various representations of the individual’s genotype allows SimBit to sustain a high performance in a wide diversity of simulation scenarios. SimBit’s performance was extensively benchmarked in comparison to SLiM, Nemo and SFS_CODE. No single program systematically outperforms the others but SimBit is most often the highest performing program and maintains high performance in all scenarios considered.

## Introduction

Evolutionary genetics has always had a strong theoretical background. As our understanding of ecological and evolutionary processes improves, we study more and more complex processes for which mathematical modelling becomes very tedious if not impossible. For such processes, only numerical simulations can allow us to perform realistic modelling. In fact, to my knowledge, the first work in computational biology has been conducted by one of the fathers of population genetics, Ronald A. Fisher (1950).

We are today in an uncanny valley in which we are almost able to perform realistic genome-wide simulations of populations but not quite yet. Individual-based simulations are used to investigate phenomena in evolutionary biology and ecology (e.g. Gilbert et al., 2017; Yeaman & Whitlock, 2011), to question conservation scenarios (e.g. Cowley, 2008; Halls & Welcomme, 2004) and are also used in statistical settings such as with Approximate Bayesian Computation (reviewed in Beaumont, 2010) or with a machine learning algorithm (e.g. Schrider & Kern, 2018). However, such technics are often computationally very expensive and it can take a lot of time to parametrize these simulations. As a consequence, many studies limit forward simulations to unrealistically low number of individuals or loci.

Writing an algorithm to make efficient individual-based simulations is no easy task, and most authors therefore rely on existing, flexible simulation programs. It is often difficult, however, to choose a simulation program. There are no objective ways to compare and express how user-friendly a program is. Also, different program packages have drastically different performance for different simulation scenarios. Learning how to use a new program can be a lengthy and difficult task, therefore many users just use the program they already know or just pick one program that is able to perform the simulations they need without questioning its performance. However, as shown below, even under simple scenarios, a given program can be hundreds or thousands of times slower than another one which will drastically affect the feasibility, or level of replication, of a study.

Here, I present SimBit, a general purpose forward-in-time population genetics simulator written in C++. SimBit has been designed to have a high performance for a wide variety of simulation scenarios. SimBit does so by using diverse representations of the genetic architecture for different simulation scenarios. As a user of Nemo (Guillaume & Rougemont, 2006), SFS_CODE (Hernandez, 2008; Hernandez & Uricchio, 2015) and SLiM (Haller et al., 2019; Haller & Messer, 2017, 2019), I gathered my experience to make SimBit a program that offers a fast learning curve to new users. With a simple set of commands that are very flexible, users can quickly simulate a great diversity of scenarios. SimBit can simulate a wide variety of selection scenarios (any selection coefficient and dominance coefficient at any locus, any epistatic interaction with any number of loci, any spatial and temporal changes of selection scenarios, etc.), demographic scenarios (any number of discrete patches with specific migration scenario, hard vs. soft selection, changes in patch size depending on fecundity, exponential vs logistic growth, gametic or zygotic dispersion, etc.), mating systems (any cloning rate and selfing rate, hermaphrodites or males and females), different types of representation of the genetic architecture (bi-allelic loci, QTLs, etc.) and SimBit has a great diversity of tools to manipulate simulations and gather output. Finally, SimBit comes with a R wrapper that is very handy for managing the creation of numerous input commands. This article aims at presenting the general working of SimBit and compares its performance to other similar programs. For detailed information about how to use SimBit, please consult the manual.

### Demography and species ecologies

In the current version, SimBit assumes non-overlapping generations (although different species can have different generation times), diploidy (although one can mimic haploidy), and assumes discrete patches (although patches can be made arbitrarily small, essentially mimicking continuous space). Outside of these three assumptions, SimBit can simulate very diverse types of scenarios. SimBit can simulate any number of patches with any migration matrix, carrying capacity, variation of the patch size from the carrying capacity based on realized fecundity with exponential or logistic growth model (the growth model can be set for each patch independently; see more on that below). Each patch can be initialized at the desired size and all of the above parameters can vary over time. Dispersal can happen at the gametic or at the zygotic phase and may be a function of the patch mean fitness (hard versus soft selection). SimBit can also simulate multiple species and their ecological interactions as explained below.

SimBit can simulate realistic changes in population in response to patch mean fitnesses. Let’s denote at time *t* the expected number of offspring of a species *s* produced in patch *p* as 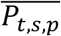. Let’s also denote the patch growth rate *r*_*t,s,p*_ = *f* Σ*w*_*i*_ as the product of *f*, the theoretical maximum fecundity of an individual having a (relative) fitness of 1.0 (set by the user), and Σ*w*_*i*_, the sum of finesses in this patch. If the user allows the patch size to vary from the carrying capacity of this species and that at time *t*, in patch *p*, for species *s*, the carrying capacity is set to K_*t,s,p*_ then the expected number of offspring produced is 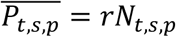 for the exponential model and 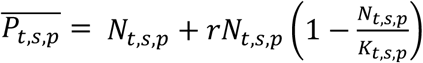 for the logistic model, where *N*_*t,s,p*_ is the size of the patch *p* of species *s* at time *t*. The actual number of offspring produced, *p*_*t,s,p*_ can then either be set deterministically 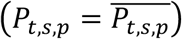 or stochastically 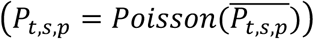). With more than one patch, these offspring produced are then spread out through migration. With a single patch (or in absence of immigration and emigration for the patch *p*), *N*_*t*+1,*s,p*_ is simply set to *p*_*t,s,p*_.

Into the above framework, we can add the fact that different species can affect each other’s through their ecological relationships. This can be achieved through a “competition matrix” that implements a Lotka-Volterra model of competition and/or through a “predation matrix” that implements a consumer-resource model (or predator-prey model) with a linear rate of resource consumption (introduction to these models in Otto & Day, 2007; discrete-time example of a predator-prey model in Çelik & Duman, 2009). Let α_i,s_ be an element of the “competition matrix” describing the competitive effect of species *i* on focal species *s*. The expected number of offspring produced is then given by 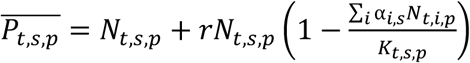. Note that competitive effects can only be set on species and on patches having logistic growth. Let β_i,s_ be an element of the “predation matrix” describing the effect of species *i* on species *s*. The predation effect is added to the expected number of offspring produced 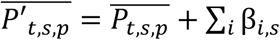. In this last equation, I assumed that all effects β_i,s_ are independent of the patch sizes of both the causal and recipient species but in practice a user can specify for each β_i,s_ whether the effect should be multiplied by the causal species patch size (*N*_*t,i,p*_), by the recipient species patch size (*N*_*t,s,p*_) or by both. SimBit enforces that all the diagonal values α_*s,s*_ = 1.0 and that all the diagonal values β_*s,s*_ = 0.0. SimBit can also allow the patch size to overshoot the carrying capacity *K*_*t,s,p*_ up to an arbitrary large value allowing for oscillating or chaotic changes in patch sizes.

### Mating system

SimBit can simulate hermaphrodites or males and females with an arbitrary sex-ratio. At every reproduction event, an organism will be cloned with probability *C* and self with probability *S*. By default, the cloning rate is set at 0.0 and the selfing rate is set at 1/2N (Wright-Fisher model), but these can be set by the user.

### Types of loci and selection

Different programs use different representations of the genetic variation. For example, Nemo represents an individual’s haplotype with an array in which the n^th^ element of the array indicates the allelic value for the n^th^ locus. In SLiM, each individual’s haplotype is represented with a container of mutations (where each mutation is an object that stores its position and other associated features as attributes). In SFS_CODE, a haplotype is represented with a linked list of mutations. These different representations of the genetic variation have important consequences for the performance of the software package. Nemo’s technique is expected to perform well at high genetic diversity per locus, while SLiM and SFS_CODE are expected to perform better at low genetic diversity per locus. Nemo also has QTLs and SLiM can mimick QTLs through Eidos (the programming language used to parameterize SLiM simulations). These different representations also have consequences on the flexibility and performance of a program.

SimBit implements five different representations of the genetic variation called T1, T2, T3, T4 and T5. I refer to these representations as types of loci. T1, T4 and T5 types of locus represent binary loci. SimBit has multiple representations of binary loci in order to sustain flexibility and high performance over a wide range of genetic diversity and of simulation scenarios. T2 type of locus represents blocks that count mutations, T3 type of locus represent QTLs and all three types. More information on these five types of representations is below. Loci of different types are integrated on the same recombination map. The recombination rate can be specified between any pair of adjacent loci (whether the two loci are of the same type or not) with any number of chromosomes. Mutation rates can also be set independently for each locus.

For a number of types of loci (see below), SimBit can make use of an assumption about the selection scenario that can provide substantial improvement in run time. I call this assumption the “multiplicative fitness” assumption. The multiplicative fitness assumption assumes 1) multiplicative fitness interactions among loci and 2) that the fitnesses of the three possible genotypes at a given locus are 1, 1–*s* and (1–*s*)^2^. When a user makes this assumption, SimBit partitions a haplotype into blocks and computes the fitness value for each block. If, during reproduction, no recombination events happen within a given block, then SimBit will not need to recompute the fitness for this specific block as the fitness of the block can simply be multiplied by the fitness of the same block on the other haplotype. By default, SimBit attempts to estimate the optimal size of these blocks, but a user can also explicitly specify the position and location of each block. This technique yields substantial performance improvement in terms of CPU time especially when the recombination rate within blocks is relatively low (see ‘Performance’ section below). Therefore, unless the exact dominance relationship is of central importance, it is generally recommended to make use of this assumption.

The genetic architecture can be set independently for each species and all the selection scenarios presented below can be set differentially for each species, habitat and time. By default, all of the patches belong to the same habitat, but a user can assign each patch to a specific habitat and all the selection pressures described below (including epistasis) can be specified for each habitat independently. Also, selection can be applied on viability and/or on fertility.

#### T1 loci

T1 loci track binary variables (e.g., mutated vs wildtype). SimBit has in memory for each haplotype an array of bits of the length of the number of T1 loci simulated. The *n*^th^ bit indicates whether the *n*^th^ T1 locus of this haplotype is mutated or not. As such, T1 loci are somewhat similar to Nemo’s genetic representation. T1 loci have high performance for simulations with very high per locus genetic diversity.

Selection scenarios on T1 loci are extremely flexible. A user can set the fitness values of each of the three possible genotypes at each locus allowing for any kind of dominance scenario including overdominance and underdominance. Any epistatic interactions between any number of loci can also be specified. A user can also use the assumption of “multiplicative fitness” on T1 loci.

#### T2 loci

T2 loci are meant to represent aggregate blocks of loci, and, SimBit counts the number of mutations happening in this block. This type should be used only when 1) the genetic diversity per T2 locus is very high, 2) when performance is a major concern, 3) the user is satisfied with the limited selection scenario it can model, and 4) a simple count of the number of mutations happening per T2 locus for each haplotype is a sufficient output. Selection on T2 is forced to have multiplicative effect among haplotypes (therefore T2 loci always use the assumption of “multiplicative fitness”).

#### T3 loci

T3 loci are quantitative trait loci (QTL) and code for an *n*-dimensional phenotype. The user can set the phenotypic effect of each T3 locus on each of the *n* axes of the phenotype, and these phenotypic effects can also depend on the environment in order to simulate a plastic response. A user can also add random developmental noise (drawn from a Gaussian distribution) in the production of a phenotype in order to reduce heritability. For T3 loci, the user can define a fitness landscape, where an individual’s fitness is given by its phenotype.

#### T4 loci

For T4 loci, SimBit computes the coalescent tree of the population over time and adds the mutations onto the tree when the user asks for output. As a consequence, T4 loci are necessarily neutral. T4 loci are inspired from Kelleher et al. (2018) and the method has already been implemented in SLiM (Haller et al., 2019). Tree recording technics can be very promising when dealing with lots of highly linked neutral loci. This technic allows a forward-in-time simulator to perform equally than backward-in-time simulators for some extreme simulation scenarios while retaining many of the advantages of forward-in-time simulations such as simulating selection at other loci (Haller et al., 2019).

#### T5 loci

T5 loci are very similar to T1 loci (two simulations with the same random seed differing only by the fact that one uses T1 loci and the other uses T5 loci will produce the same output). For each haplotype, SimBit has a dynamic sorted array with the position of each T5 locus that is mutated. As such T5 loci are somewhat similar to how SLiM keeps track of its genetic architecture. With high genetic diversity SimBit therefore tracks a lot of mutated loci, while with low genetic diversity SimBit tracks few mutated loci. For this reason, T5 loci tend to perform better than T1 loci for moderate to low genetic diversity per locus.

Behind the scene, SimBit will track separately T5 loci that are under selection and T5 loci that are neutral for improved performance. SimBit can also compress T5 loci (either the neutral ones and/or the selected ones) information in memory. Compression reduces the RAM usage by up to a factor of 2 and can increase or decrease CPU time depending on the simulation scenario. By default, SimBit makes this compression on the neutral T5 loci only and only when it is certain it will improve performance. For advanced users, it is also possible to ask SimBit to invert the meaning of some loci depending on their frequencies. For example, if the locus 23 is fixed or quasi-fixed, haplotypes would track this 23^rd^ locus only if they carry the non-mutated allele.

With T5 loci, one can specify the fitness values of the heterozygote and double mutants’ genotypes only allowing for all types of dominance including overdominance and underdominance. Just as on T1 loci (and T2 loci), a user can take advantage of the assumption of “multiplicative fitness”.

### Initialization

Several options exist in SimBit to initialize and reset the genome of existing individuals. The patch size as well as the genetic diversity for each locus can be set at initialization. A user can then perform any mutation desired at predefined times with the option --resetGenetics. To ease user interface, SimBit also allows the user to define “individual types” (via option --individualTypes). Those individual types can then be used to either initialize a population or to insert (or replace) new individuals into any patch at arbitrary moments (also via option --resetGenetics). One can, for example, create individual types belonging to large hypothetical patches and simulate immigration from these hypothetical patches by just introducing these individual types into the focal patch. This speeds up simulations as SimBit does not explicitly simulate these large source patches. It is also possible to start a simulation from the individuals of a previous simulation that have been saved in binary files. Binary files are particularly useful to 1) avoid simulating a burn-in multiple times, 2) resume a simulation from an intermediate timepoint, and 3) save the entire population in a compact format to extract specific summary statistics later on.

### Outputs

Outputs are often very limiting factors for population genetic simulators (Hoban et al., 2012). SimBit can produce 30 different types of outputs (which can be sampled at any number of generations throughout the simulation). These outputs include, but are not limited to, entire genotypes of each individual in the metapopulation, allele frequencies, *F*_*ST*_, VCF files, fitness (specifying fitness for each type of locus), patch sizes, extinction times of the different species, the whole genealogy between two specified generations, binary files of the entire population (that can be reused for future simulations or simply to extract summary statistics later on). Many of these outputs can be restricted on a specified subset of loci. SimBit can also simulate sequencing errors before producing the outputs to make results easier to compare to empirical data.

### User interface

SimBit reads options either directly from the command line or via an input file. An important goal of SimBit is to have a user interface that takes input that is readable and in a very simple format to give the users a good understanding of what they are simulating and offer very explicit error messages when input is nonsense. SimBit recognizes specific options as they are proceeded by a double dash (‘--‘). For example, ‘--patchCapacity unif 1e4‘ indicates that the carrying capacity is uniform (keyword ‘unif‘) for all patches and is set to 10,000. The ordering of these options does not matter. SimBit also provides a number of macros that are mainly inspired from R functions. These inputs can be read either directly from the command line or from a file. SimBit also comes with an R wrapper.

In order to be fast and easy to learn, SimBit provides many functionalities with a relatively small number of options. It achieves this by having most options being specific to a generation, a habitat and/or a species and uses specific markers, @G, @H and @S to input information that are generation-specific, habitat-specific and species-species, respectively. For example, the entry -- N @G0 unif 100 @G5e3 unif 1000 asks for the carrying capacity of all patches to be uniformly (keyword `unif`) set to 100 from generation 0 to generation 4999 and then set to 1000 until the end of the simulation. Also, most options come with a diversity of modes of data entry. For example, for the migration scenario, a user can indicate the whole dispersal matrix or can simply specify an island model, a linear stepping stone model or a Gaussian dispersal kernel. Below, I benchmark SimBit in comparison to other softwares. Examples of command line inputs to SimBit for these simulations which results are shown on figures 1, 2, S1, S2, S3 and S4 as well as for the simulations of figure 3 are found in appendix A. Here is an example of a input file used for this benchmark. Please see manual for more information.

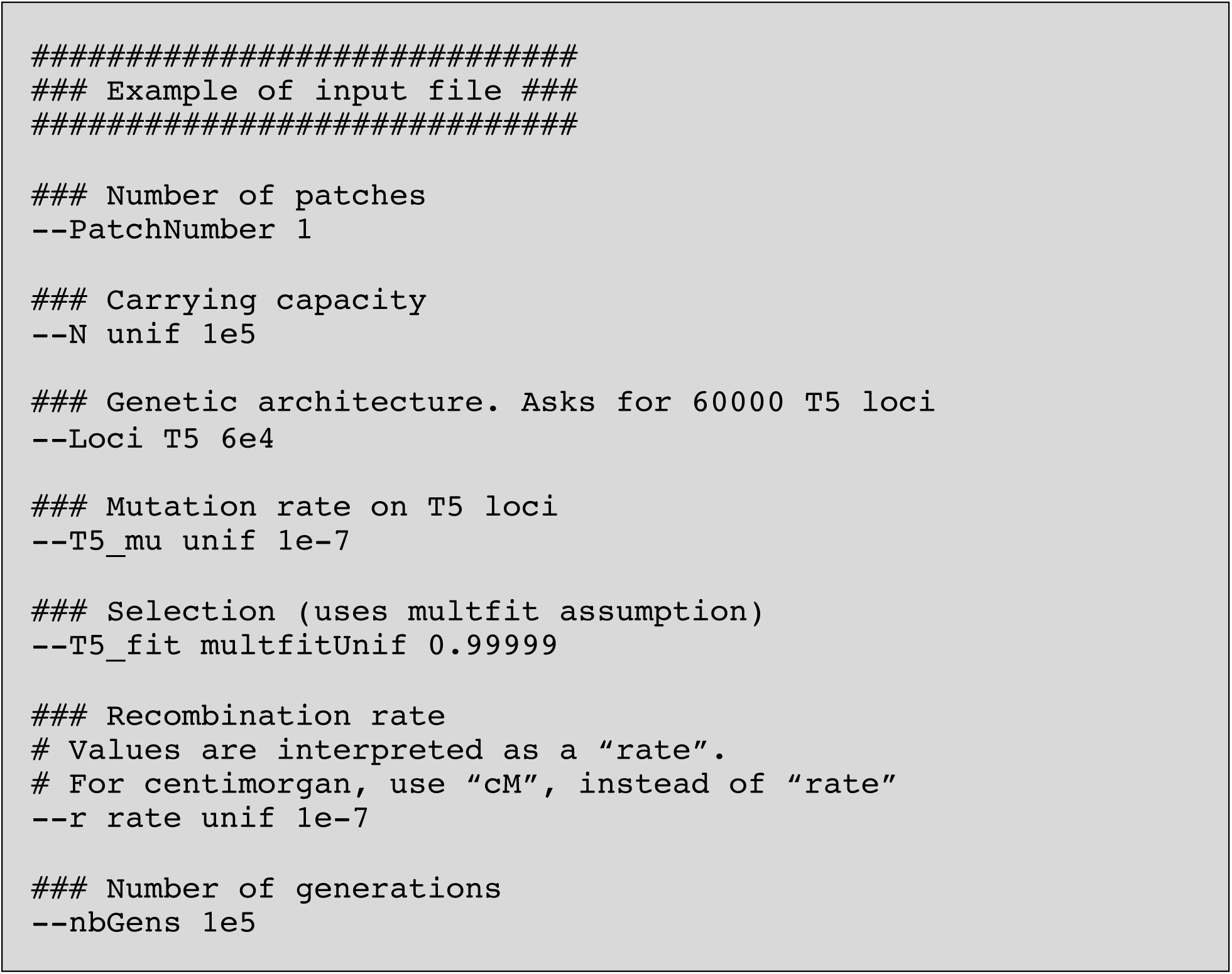

**Figure 1:**
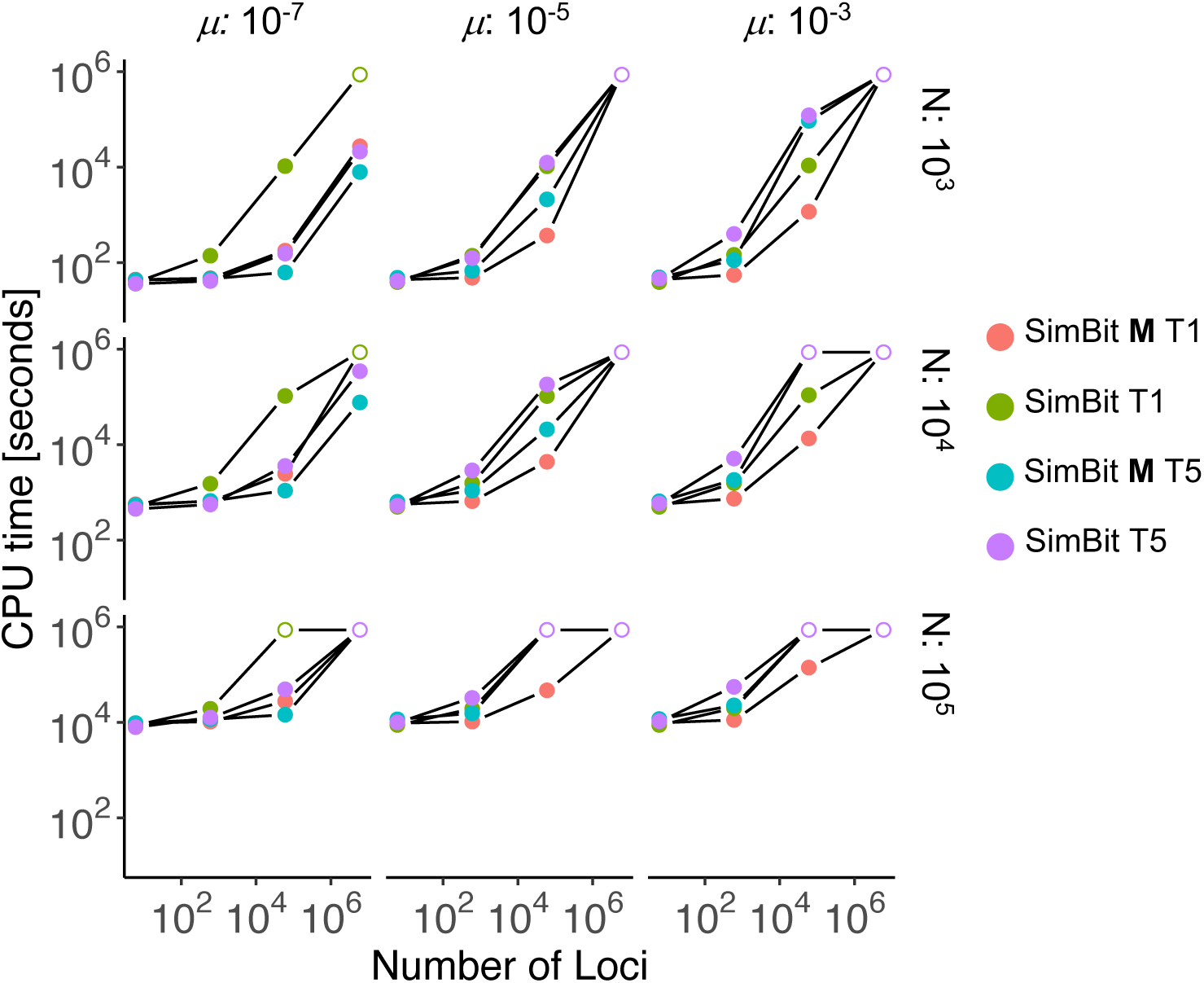
Comparison of computational time among the four different ways to simulate the same evolutionary scenario using SimBit. Results here are only for a subset of parameters (excluding N=100, N=10^6^ and all scenarios where the recombination rate among adjacent loci differs from 10^−7^). Other scenarios are in figure S1. Comparisons of memory usage (max Resident Set Size) are found in figure S2. Simulations that exceeded 10 days (240 hours) of simulation time or 20GB of memory were killed and are reported below with an empty dot at 240 hours (8.64 × 10^5^ second). The bold **M** signifies the usage of the assumption of multiplicative fitness.

**Figure 2:**
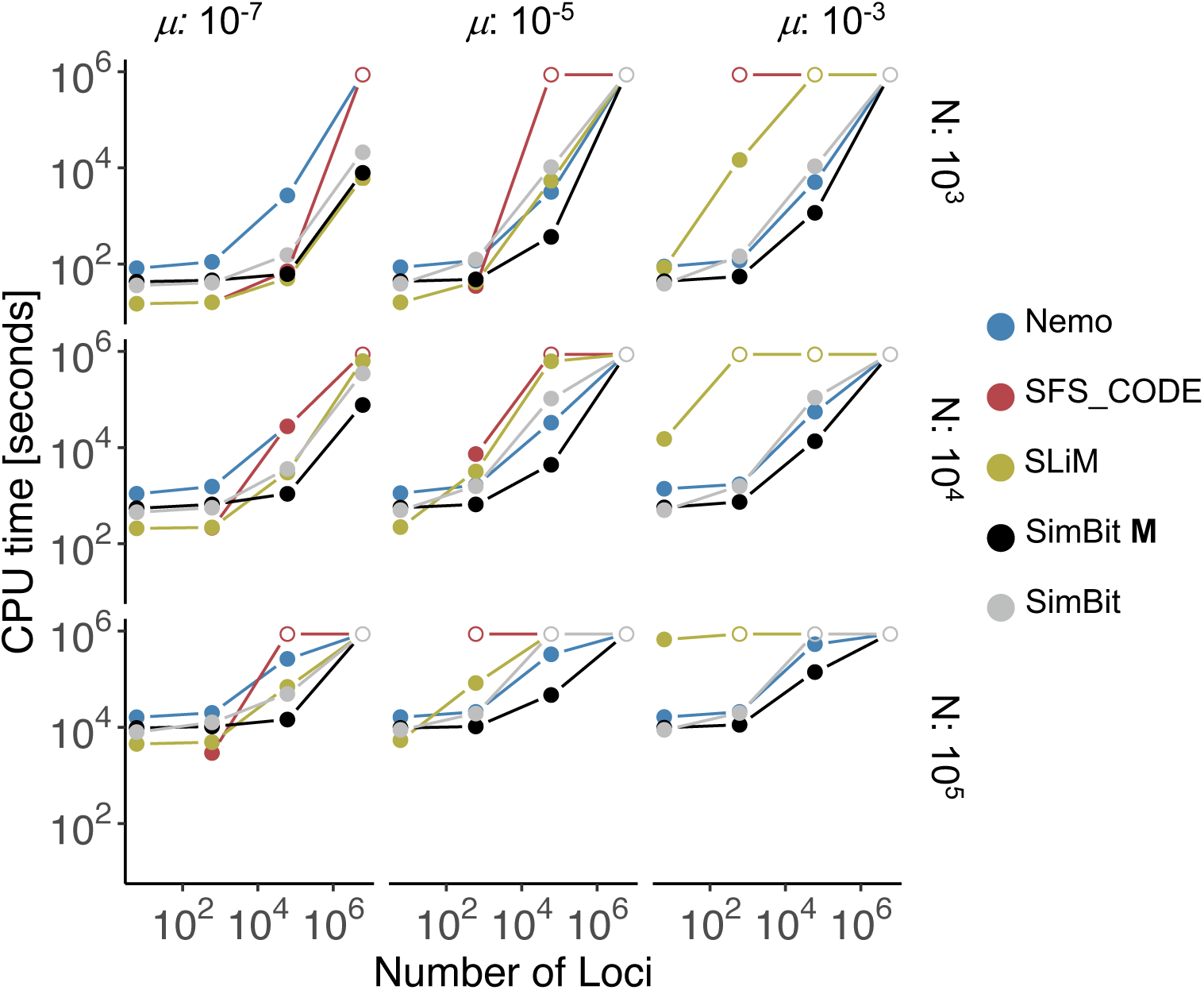
Comparison of computational time among the four different simulation programs Nemo, SFS_CODE, SLiM and SimBit. For SimBit, two lines are displayed showing the best performing between T1 and T5 loci from figure 1, once taking advantage of the assumption of multiplicative fitness, once without taking advantage of this assumption. For comparison, SLiM and Nemo are unable to take advantage of this assumption while SFS_CODE is forced to make this assumption. Other scenarios are in figure S3. Comparisons of memory usage (max Resident Set Size) are found in figure S4. See figure 1 for more details.

**Figure 3:**
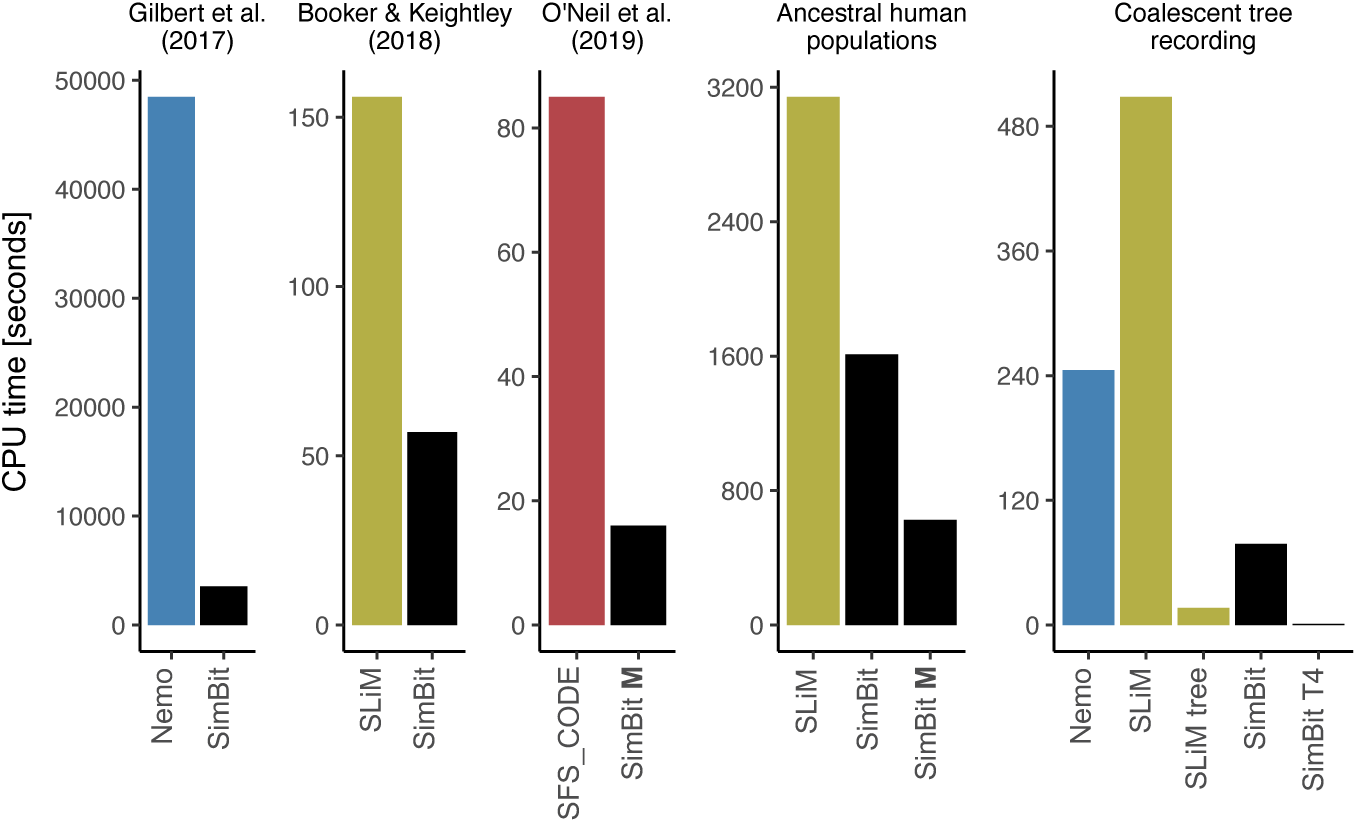
Comparison of CPU time among the four programs to reproduce simulations inspired from three recent papers as well as for a neutral simulation scenario with extreme parameters chosen to highlight the possible advantage of T4 loci (Tree recording). The bold **M** signifies the use of the assumption of multiplicative fitness. SFS_CODE simulation from the “Neutral simulation example” as well as both SFS_CODE and Nemo simulations from the “Human ancestral populations” were purposely killed after overpassing 50 times SimBit’s CPU time for the same simulation.

SimBit also comes with an R wrapper that is particularly useful for building numerous input simulations. Without going into explaining the detail working of the wrapper, let’s consider a complete example of code that will test how different migration rates and number of patches in an island model affect *F*_*ST*_. The first step is to create a grid of parameters (a “data.frame”), where each row contains information for a single simulation. We will use a full factorial design with three distinct migration rates and seven distinct number of patches. We will run 20 replicates for each of these 3×7=21 combinations resulting in a grid of parameters of 420 rows. The argument “outputFilePrefix” sets a column called “outputFile” with the prefix given followed by the row number. This column will be used to set the where outputs should be directed.

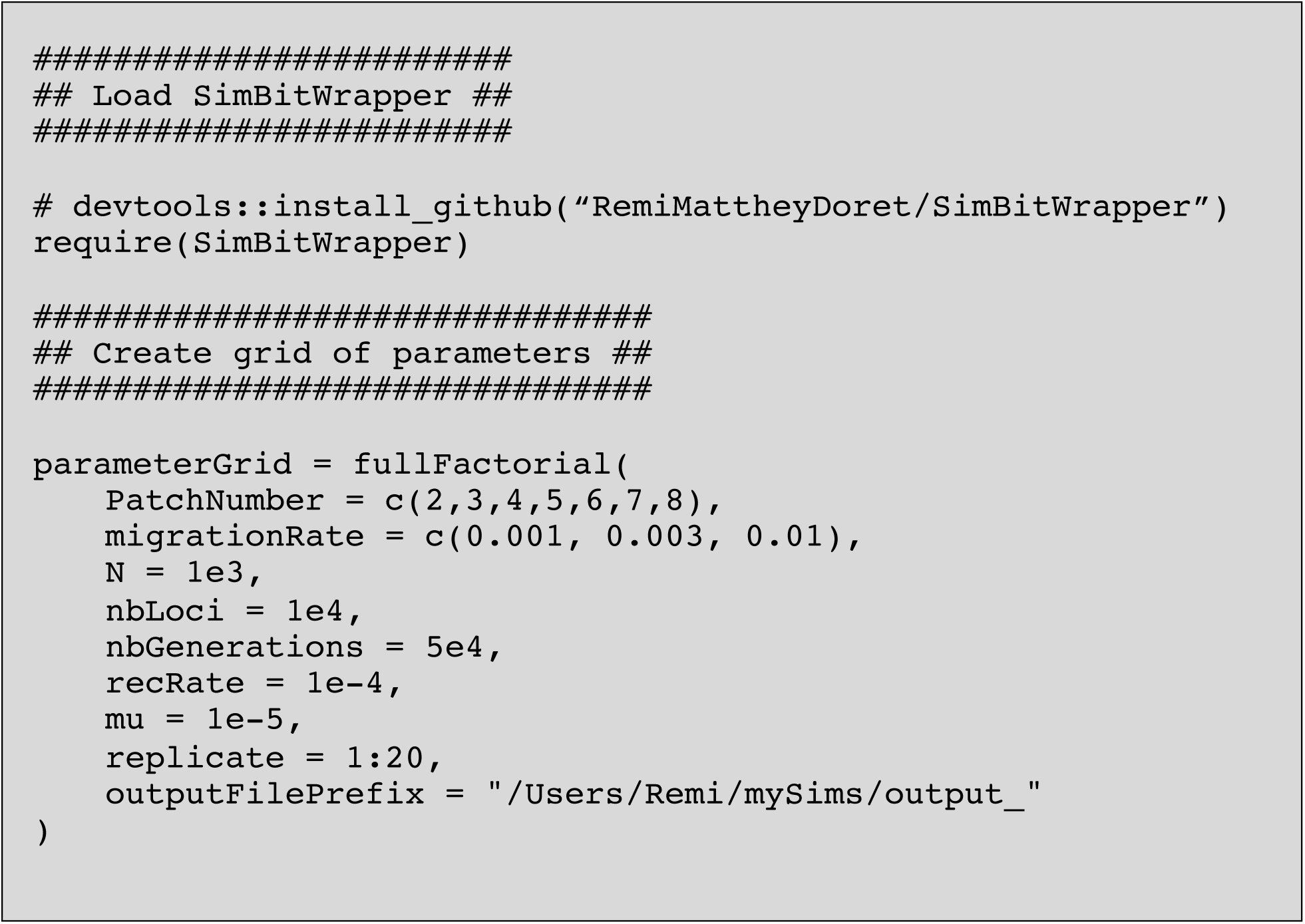

The second step is to loop through the rows of the parameter grid in order to run the simulations (or to create the input file to run them later on). For this, we use the function `GetParameterGridData`, which, for each column of the grid of parameters, sets a variable with name equal to the column name and value equal to the value of this column at the specified row of the specified parameter grid given in input.

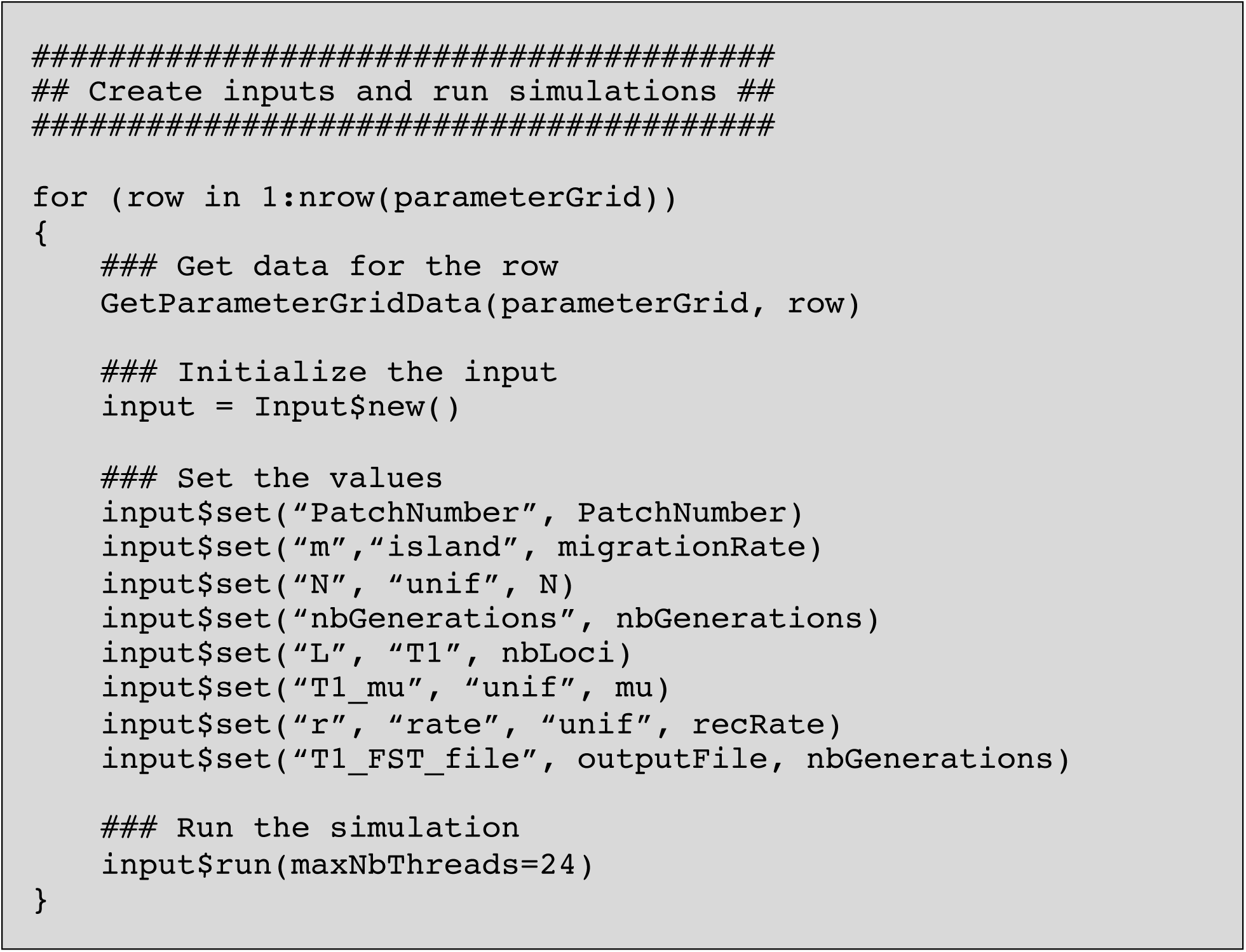

The argument `maxNbThreads` is an easy way to parallelize the simulations. `maxNbThreads`=`24` does not mean that a given simulation will use 24 threads (each simulation takes one thread) but that the `run` method will start 24 simulations in the background and will then wait that one of them finishes before starting a 25^th^ simulation. Please see manual for further information about the `run` method. It is sometimes more practical to print the input command into a file either and run the simulations from the shell at a later time. This can be achieved with `input$print(“/path/to/input.txt”)`. Finally, the last step is to gather the outputs and graph the results. In order to gather the outputs, we use the function `gatherData`. This function uses a number of optional parameters (see manual) but default parameters work fine for our simple example.

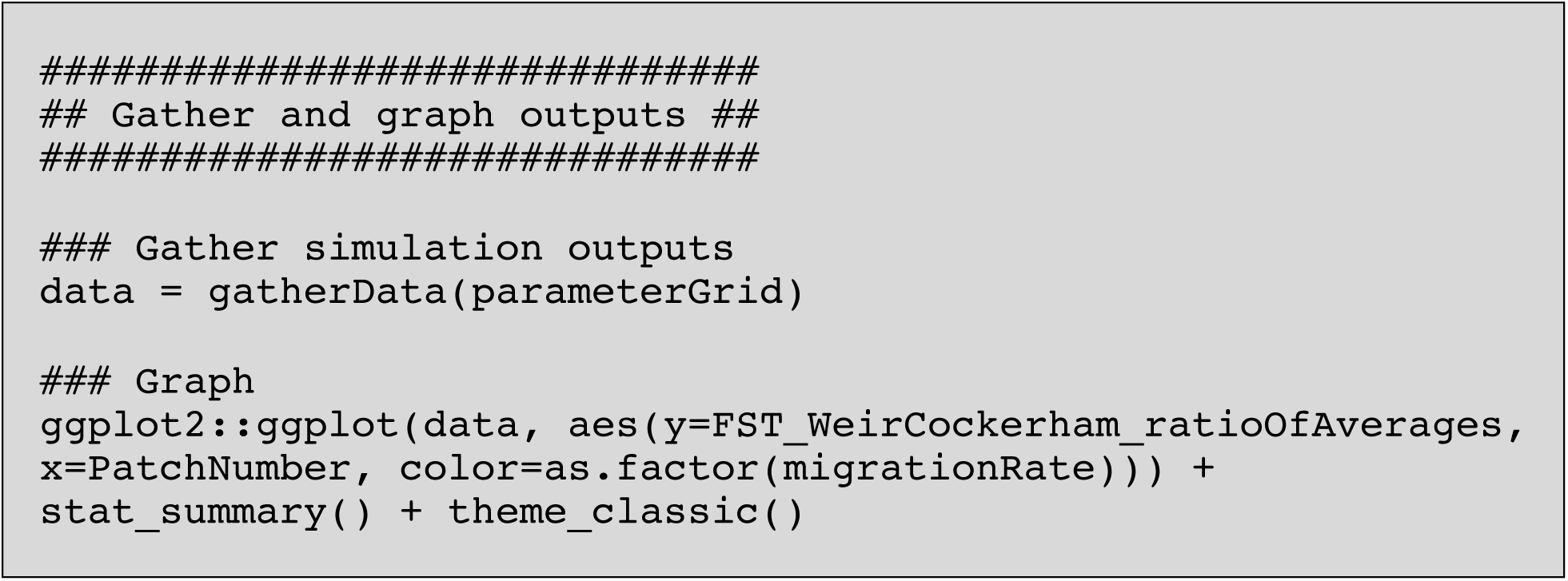

In this simple example, the entire study (defining the parameters, creating the inputs, running the simulations, gathering and graphing the results) takes 16 lines of code (16 expressions; including loading packages, excluding the curly braces; and it could be reduced to 7 lines only)! The column “FST_WeirCockerham_ratioOfAverages” used for plotting corresponds to Weir & Cockerham (1984) estimator of *F*_*ST*_. The resulting graph is displayed in figure S5 on which is added the theoretical expected *F*_*ST*_ values from Charlesworth (1998) for comparison.

### Program comparison – Performance

It is often hard for a user to know which program to use for a given study. Indeed, few articles compare program’s features (but see Hoban, 2014, who compares software flexibility), and when authors publish a new program, they do not always compare its performance to other similar programs (but see performance comparisons between SLiM, SFS_CODE and fwdpp in Haller & Messer, 2017).

In this article, I compared performance of SimBit to three forward-in-time programs; SFS_CODE (Hernandez, 2008; Hernandez & Uricchio, 2015), SLiM (Haller et al., 2019; Haller & Messer, 2017, 2019; Messer, 2013) and Nemo (Guillaume & Rougemont, 2006). I chose these three programs because they are all forward-in-time simulation platforms, they can all simulate selection, they are all popular (392 citations among the articles announcing SLiM, SLiM2, SLiM3 and the implementation of tree recording sequences in SLiM; 127 citations for Nemo; 216 citations for SFS_CODE; as of 23rd April 2020 on Google Scholar) and are generally considered the highest performing software available.

SimBit contains a number of options that are meant to refine its performance (see section “Performance options” in the manual). In practice though, most users will probably only need to choose the type of loci to simulate, and SimBit will do a decent job to figure out how best to simulate it. In order to best represent the performance that a new user ought to expect from SimBit, however, all simulation performances (CPU time and memory usage) presented below are made with the default parameters of SimBit.

In order to compare program performance, I ran basic simulations with a single Wright-Fisher population, uniform mutation rate and a uniform recombination rate. All loci experienced a selection coefficient of *s*=0.00001 and *h*=0.5. Low selection coefficients were chosen to 1) prevent any software from throwing an error stating that it might suffer from round-off errors caused by low mean fitness and 2) reduce the effects of assuming multiplicative fitness among haplotypes on the simulated scenario (fitness differences between simulations that take advantage of the assumption of multiplicative fitness and the ones that do not is of the order of 10^−11^). Note that while SimBit can take advantage of this assumption of multiplicative fitness on demand, SFS_CODE is forced to make this assumption and Nemo and SLiM cannot take advantage of this assumption. I varied the mutation rate (taking values 10^−7^, 10^−5^ and 10^−3^ per locus), the recombination rate (taking values 0, 10^−9^ and 10^−7^ and 10^−5^ per adjacent locus), the carrying capacity (taking values 10^2^, 10^3^, 10^4^, 10^5^ and 10^6^ diploid individuals), and the number of loci (taking values 6, 6×10^2^, 6×10^4^ and 6×10^6^) in a full factorial design. All simulations ran for 10,000 generations. I ran these simulations with Nemo (version 2.3.46), SLiM (version 3.1), SFS_CODE (version 20150910) and SimBit (version 4.11.0). Because using Nemo’s full potential is not trivial, for Nemo, the input files used for these benchmarks were directly created by Frederic Guillaume. In order to compare the behaviour of different types of loci and selection scenarios in SimBit, I ran all simulations four times in SimBit with T1 and T5 types of loci with and without making use of the assumption of multiplicative fitness among haplotypes. CPU time and peak in Resident Set Size (RSS; memory) usage are reported. Simulations that exceeded 10 days (240 hours) of simulation time or 20GB of memory usage were killed and are reported below with a dot at 240 hours (8.64 × 10^5^ seconds in the units used on the figures) and at 20GB (2 × 10^7^ kb in the units used on the figures). All these simulations were run on an Intel Xeon X5650 processor and codes were compiled with gcc-4.8.2rev203690. I ensured that the number of SNPs were not significantly different between all four programs for three of the simulation scenarios benchmarked.

For brevity and because changing the recombination rate has very little effect on the results (only SFS_CODE appears to significantly slow down with higher recombination rates), I am showing only the recombination rate 10^−7^ and only the carrying capacities 10^3^, 10^4^ and, 10^5^ in the main figures. The other benchmarks are found in supplementary material. Figure 1 compares the CPU time among SimBit simulations (T1 vs. T5 and with vs. without taking advantage of the assumption of multiplicative fitness among haplotypes) for a subset of scenarios. Figure S1 and S2 compare, respectively, the CPU time and the memory usage among SimBit simulations for all scenarios. Figure 2 compares CPU time among Nemo, SLiM, SFS_CODE and SimBit for a subset of scenarios. Figure S3 and S4 compare, respectively, the CPU time and the memory usage among Nemo, SLiM, SFS_CODE and SimBit.

As expected, T1 loci perform best at high per locus genetic diversity, while T5 loci perform best at moderate to low per locus genetic diversity (figure 1). This is because with T5 loci, SimBit tracks the mutated loci, while with T1 loci, SimBit tracks every locus whether mutated or not (see above section “Representations of the genetic architecture”).

Simulations taking advantage of the assumption of multiplicative fitness generally performed better. This advantage decreases as recombination gets higher. For the range of recombination rates explored (up to 10^−5^ among adjacent loci), simulations taking advantage of the assumption of multiplicative fitness always outperformed the simulations that did not make this assumption. The reason why recombination rate matters for performance is because, as explained in section “Types of loci and selection”, SimBit needs to recompute fitness for a fitness block only if a recombination event happens within this block when using the multiplicative fitness assumption.

Comparisons between different programs highlight that there is no one program that always performs best (figure 2; figure S3). However, unlike all other software tested, SimBit perform highly in all simulation scenarios considered. SFS_CODE’s CPU time and peak RSS increases exponentially with increase in mutation rate and population size (see also simulations performed by the Ryan Hernandez on SFS_CODE websites; sfscode.sourceforge.net/SFS_CODE/Performance.htlm). Hence, SFS_CODE performs well for simulations that have very low genetic diversity, but it quickly becomes very slow as genetic diversity increases.

Nemo is most competitive when there is high genetic diversity per locus (high mutation rate and high population size). This was expected because Nemo tracks every single locus for each haplotype whether or not it is mutated. In fact, with high genetic diversity, Nemo sometimes runs in less time than SimBit when SimBit did not take advantage of the multiplicative fitness assumptions (the grey dots in figures 2 and S3). Nemo never outperformed SimBit in terms of memory usage though (Figure S4) or in terms of CPU time when SimBit takes advantage of the multiplicative assumption.

SLiM, just like SFS_CODE, performs best at very low genetic diversity. SLiM computational time is however not as exponential as SFS_CODE, which makes SLiM fast for a wider range of simulation scenarios. SLiM tends to perform better than SimBit when there is little genetic diversity, while SimBit tends to perform better when there is moderate to high genetic diversity. In general, performance comparison in terms of memory usage (figures S2, S4) mirrors well the performance comparisons in terms of CPU time (figures S1, S3).

A difference in performance is not just a question of whether a user will have to wait a little longer to get their output; often it is the difference between a research project that is feasible or not. The log scale on figures S1 and S2 (and supp. figures) might give the reader a false impression of the importance of an observed difference. Consider for example the simulation scenario where r=10^−7^, N=10^3^, *µ*=10^−7^ and 6 loci where SLiM outperforms SimBit. SLiM runs in 16 seconds while SimBit runs in 37 seconds. Let’s now consider the simulation scenario where r=10^−7^, N=10^5^, *µ*=10^−7^ and 6×10^4^ loci. SimBit (with multiplicative fitness assumption) runs in ∼4 hours, while SLiM runs in ∼19 hours, Nemo runs in more than 3 days and SFS_CODE does not manage to finish within the 10-day limit. To further consider comparisons between SLiM and SimBit as example, from figure 2, the simulation scenario where SLiM is comparably the fastest, SLiM is 2.56 times faster than SimBit; SimBit took 41 seconds while SLiM took only 16 seconds. For the simulation scenario where SimBit is comparably the fastest, SimBit is (at least) 1169 times faster than SLiM; SimBit took ∼12.3 minutes while SLiM was killed after overpassing the 240 hours walltime. These performance differences can translate into a major determinant of what can be achieved for a research project.

These very simple simulation scenarios benchmarked above might not be representative of what people really want to simulate. I therefore performed further benchmarking by comparing the performance of Nemo, SLiM, SFS_CODE and SimBit for simulations inspired by recent papers. I sampled three papers, one that performed simulations with SFS_CODE (O’Neill et al., 2019), one that performed simulations with Nemo (Gilbert et al. 2017) and one that performed simulations with SLiM (Booker & Keightley, 2018). To simplify the writing of the commands and make sure that the comparison is fair, I simplified the Booker and Keightley (2017) simulations by assuming a constant mutation rate and recombination rate and used the gamma distribution of fitness effects with a mean of 0.05 and an alpha parameter of 0.111. For the Gilbert et al. (2017) paper, the simulations have also been slightly modified from the original. The original paper’s specified a “breeding kernel” that can only run on a modified version of Nemo that is not directly published on Nemo’s official website. Hence, for the Gilbert et al. (2017) simulation, I removed the breeding_kernel and modified the size of the dispersal kernel appropriately. For simplicity (because the original input file was 390Mb large), I also used a linear stepping stone model of 8000 patches starting with the 1000 left-most patches at carrying capacity and the others empty. I made sure the expansion speed was similar among the two programs. For fairness, I compared the Nemo and SLiM that cannot take advantage of the assumption of multiplicative fitness with SimBit that does not make this assumption, while I compared SFS_CODE that is forced to make this assumption with SimBit that makes this assumption. I also performed a benchmark inspired from human genome and human ancestral demography. I simulate 500 patches of 100 individuals each in a linear stepping stone model with a migration rate to either of the two neighboring patch of 0.2. The genome contained 2×10^8^ sites with a uniform mutation rate of 2×10^−8^ and a uniform recombination rate of 10^−8^. For simplicity, all loci were under purifying selection with a constant selection coefficient of 0.0001 and a dominance coefficient of 0.5. Finally, I added a benchmark of a simple Wright-Fisher simulation scenario (N=1000, *µ*=10^−5^, 10^6^ loci, r=0; 5000 generations) without selection. Neutral loci can be tracked through a coalescent tree for both SLiM (with Tree Recording and subsequent analysis of the outputted binary file in Python) and SimBit (with T4 loci). These simulations were run on an Intel i7-8559u processor, and codes were compiled with clang-800.0.42.1.

SimBit systematically outperforms the software used in the original papers (figure 3). For the simulation inspired from human genetics and ancestral human population, SimBit outperformed SLiM whether it made use of the multiplicative fitness assumption or not. Finally, for the “Neutral simulation example”, the coalescent tree recording technique of both SLiM and SimBit vastly outperform more traditional techniques (figure 3). With “traditional techniques”, SLiM, Nemo and SimBit took 8m29s, 4m05s and 1m18s, respectively, while using coalescent tree recording methods, SLiM and SimBit only took 16.6 seconds and 1.2 seconds, respectively. Here, I only considered an extreme scenario to exemplify the possible advantage of tree recording techniques. For example, I used a recombination rate of zero. With higher recombination rates, the computational time of tree recording techniques would become slower, while it would not have much impact on the runs that did not use a tree recording technique.

## Conclusion

There is no perfect way to compare program performance, and one must always be careful when making conclusions from such a benchmark. First, the parameter space considered is, of course, finite. For example, my benchmark does not include any single-locus simulations, simulations with high selfing rates or with males and females instead of hermaphrodites, or any simulations with a very high recombination rate. Also, different programs mean different things by a locus. SFS_CODE simulate triplets of loci as a codon. This means that many mutations that are happening in SFS_CODE are synonymous mutations that don’t affect fitness. Consequently, the performance comparisons shown here are unfairly favourable to SFS_CODE compared to Nemo, SLiM and SimBit, but it would not be any fairer either to run all SFS_CODE simulations with three times as many loci. Nemo uses a byte to represent each neutral locus (but only a single bit for loci under selection) hence allowing for the representation of up to 256 possible alleles at neutral loci. SimBit on the other hand represent each locus with a single bit (whether the locus is under selection or not), hence allowing for only two possible alleles. SLiM’s mutations “stack” (no reverse mutations) at a given locus, hence simulating a pseudo infinite allele type of model (see SLiM manual on “mutation stacking” for more information; http://benhaller.com/slim/SLiM_Manual.pdf). As explained above, SimBit contains a number of performance tweaks a user can take advantage of to improve the performance above the default run mode (compression of T5 data in memory, allowing inversion of the meaning of T5 loci depending on their frequency, turning on/off the swapping of pointers for haplotypes that do not recombine or mutate during reproduction, setting manually the positions of blocks for the multiplicative fitness assumption). However, the above simulations were all performed with SimBit default values for these performance tweaks, which is somewhat unfair to SimBit.

SimBit has already been used in a number of projects. It has been used for simulations that require very high performance, simulating the effect of background selection of large stretch of DNA in structured populations (Matthey-Doret & Whitlock, 2019). SimBit has also been used for two projects on genetic rescue, one requiring habitat-specific epistatic interactions (Nietlisbach et al., forthcoming) and one requiring complex metapopulation initialization and introduction of predefined individuals during the simulation (Whitlock lab consortium, forthcoming). SimBit is under a permissive free program license and is available at https://github.com/RemiMattheyDoret/SimBit.

## Acknowledgment

Thank you to Michael C. Whitlock for his help through discussions in designing SimBit. Thank you to both Michael C. Whitlock and Kimberly J. Gilbert for helpful comments on the manuscript. Thank you to Pirmin Nietlisbach for being the main beta tester and for his advice on how to improve the user interface and the manual. Thank you also to Ben Haller and Frédéric Guillaume for their feedback on how to make a fair comparison among programs. Special thanks to Frédéric Guillaume for his help at creating the input files for Nemo and for his feedback about how to display the benchmark results in a way that is fair. Finally, thank you to ComputeCanada for the computational resources used for benchmarking.

