## Supplemental figures for "SimBit: A high performance, flexible and easy-to-use population genetic simulator"

**Figure S1:** Comparison of computational time among the four different ways to simulate the same evolutionary scenario using SimBit. Comparisons of memory usage (max Resident Set Size) are found in figure S2. See figure 1 in main text for more details.

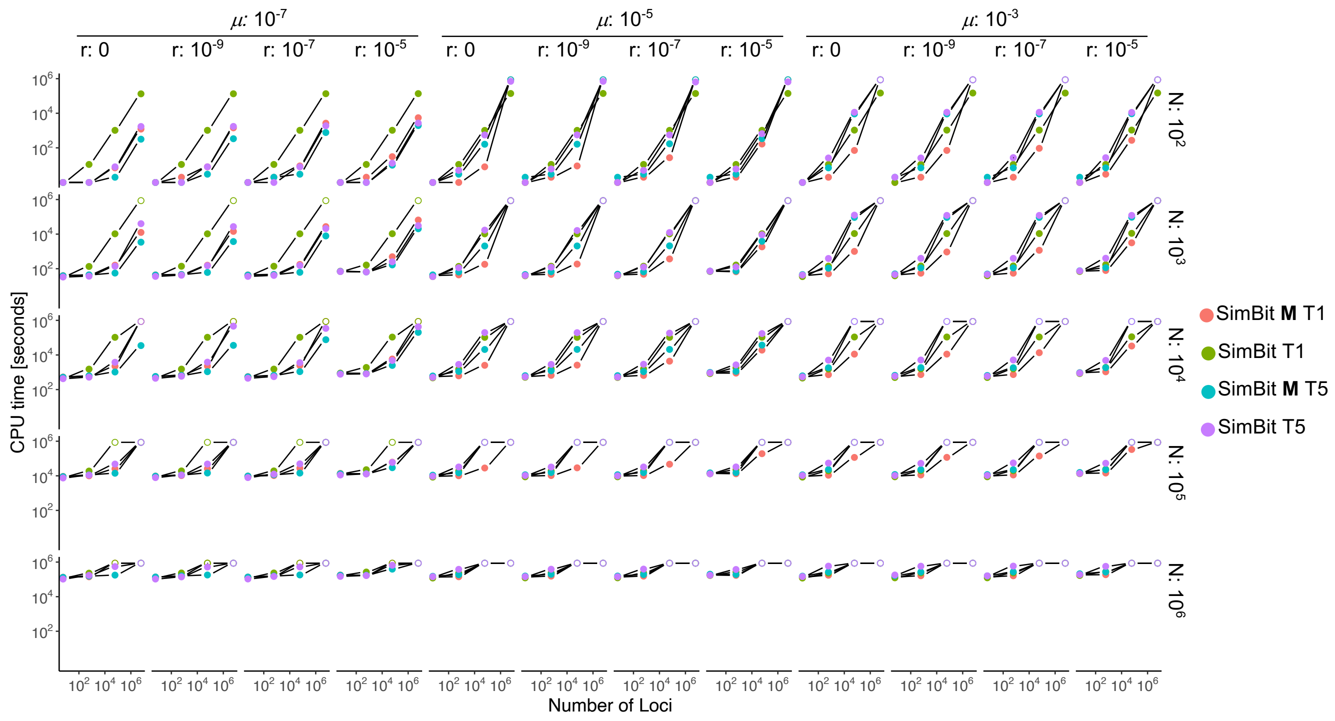

**Figure S2:** Comparison of memory usage (max Resident Set Size) among the four different ways to simulate the same evolutionary scenario using SimBit. Comparisons of CPU time are found in figure S1. See figure 1 in main text for more details

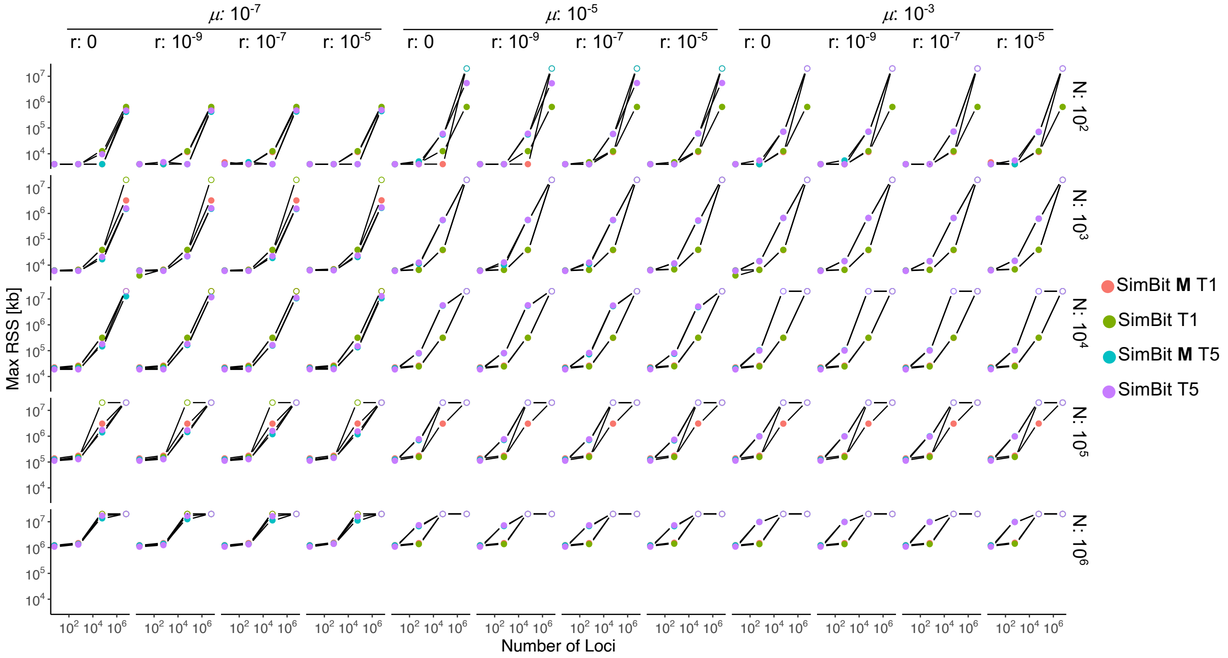

**Figure S3:** Comparison of computational time among the four different simulation programs Nemo, SFS_CODE, SLiM and SimBit. For SimBit, two lines are displayed. Both lines show the best performing between T1 and T5 loci from figure S1, once taking advantage of the assumption of multiplicative fitness, once without taking advantage of this assumption. See figure 1 in main text for more details.

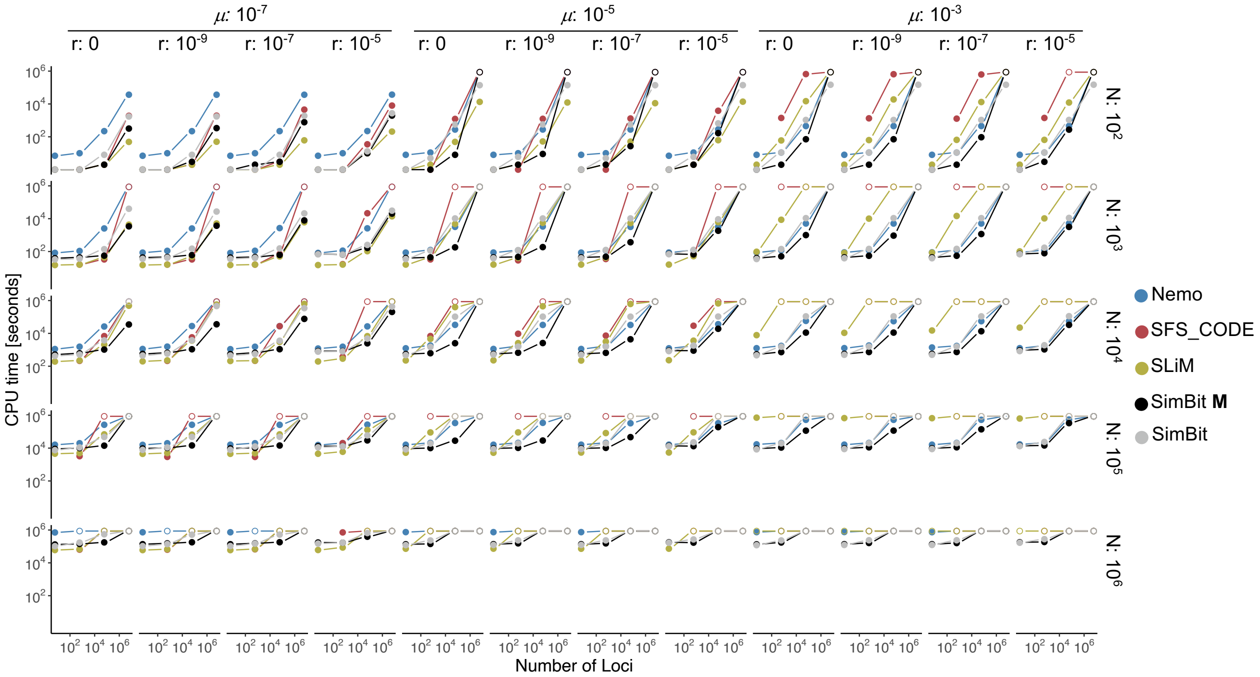

**Figure S4:** Comparison of memory usage (max Resident Set Size) among the four different simulation programs Nemo, SFS_CODE, SLiM and SimBit. For SimBit, two lines are displayed. Both lines show the best performing between T1 and T5 loci from figure S1, once taking advantage of the assumption of multiplicative fitness, once without taking advantage of this assumption. See figure 1 in main text for more details.

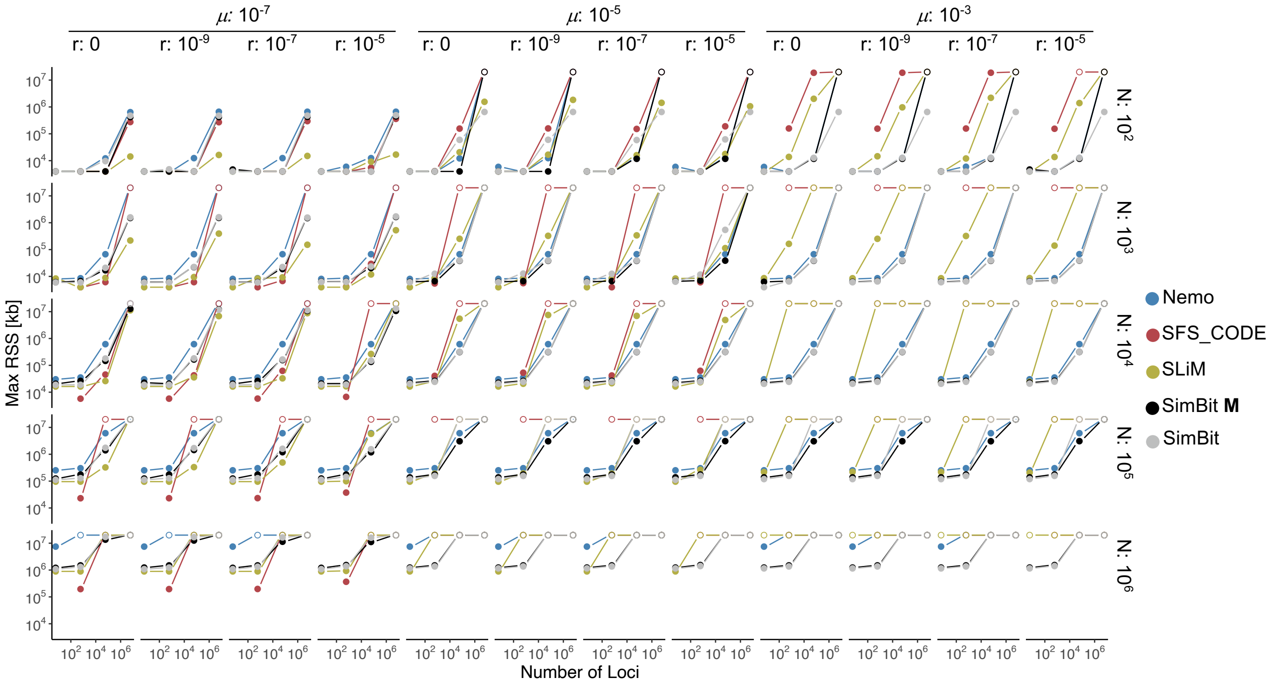

**Figure S5:** Results of the use case example presented in section "interface". The black lines represent analytical expectations from Charlesworth (1998). The $F_{ST}$ values computed with Weir and Cockerham (1984) estimator and therefore correspond to the column "FST_WeirCockerham_ratioOfAverages" in the output files. Note that the y-axis is on a logarithmic scale. Error bars represent 95% confidence interval. Most error bars are too small to be distinguished from the dot representing the mean $F_{ST}$.

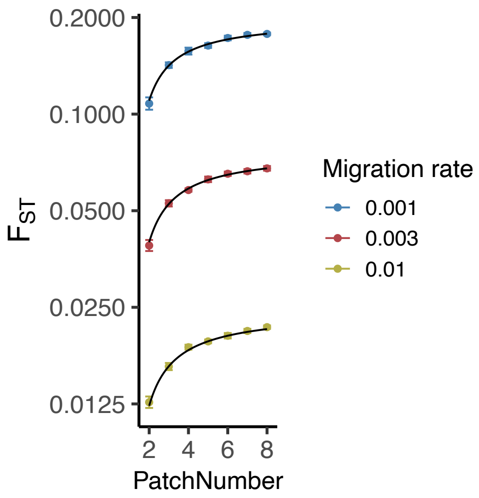
