## Appendix_A for "SimBit: A high performance, flexible and easy-to-use population genetic simulator"

Below input commands are described without using SimBit's R wrapper. Here is the input used for the Wright-Fisher simulations presented at figures 1, 2, S1, S2, S3 and S4, with 100,000 individuals, 60,000 T5 loci, a uniform recombination rate of $10^{-7}$ and a uniform mutation rate of $10^{-7}$, assuming that fitnesses among haplotypes are multiplicative.

```
SimBit --PN 1 --N unif 1e5 --L T5 6e4 --T5_mu unif 1e-7 --T5_fit multfitUnif 0.99999 --r rate unif 1e-7 --nbGens 1e5 --printProgress f
```

When written on a file, one can add comments and therefore this input might look like

```
# Number of patches (aka --PatchNumber)
--PN 1

# Carrying capacity
--N unif 1e5

# Genetic architecture (aka --Loci)
--L T5 6e4

# Mutation rate
--T5_mu unif 1e-7

# Selection (uses multfit assumption)
--T5_fit multfitUnif 0.99999

# Recombination rate
--r rate unif 1e-7

# Number of generations
--nbGens 1e5

# Avoid printing the generation counter to standard output
--printProgress f
```

'--PN 1' (could also be written '--PatchNumber 1') sets the number of patches to 1. '--N unif 1e5' indicates a uniform carrying capacity for all patch of 100,000 individuals. '--L T5 6e4' asks for 60,000 T5 loci. '--T5_mu unif 1e-7' sets a uniform mutation rate of $10^{-7}$ per locus. '--T5_fit multfitUnif 0.99999' indicates a uniform selection scenario over all loci with the fitnesses of the three being set at 1, 0.99999 and $0.99999^2$. '--r rate unif 1e-7' indicates a uniform recombination rate of $10^{-7}$ between adjacent loci. Note the keyword 'rate' indicate the unit by which the recombination rate is set. Alternatively, one could use 'cM' for centiMorgans or 'M' for Morgans. '--nbGens 1e5' (could also be written '--nbGenerations 1e5'). Finally, '--printProgress f' simply turns off ('f' is short for 'false') the printing in the standard output of the progress of the simulation.

For the simulations inspired by Gilbert et al. (2017), the input for SimBit is

```
SimBit --T1_mu unif 0.0001 --T1_fit unif 1 0.997 0.99 --PN 8000
--N unif 12 --L T1 1e3 --r rate unif 0.005 --nbGens 18000 --m
LSS 3 0.3 0.4 0.3 1 --fec 7 --InitialpatchSize A rep 12 1e3 rep
0 7e3 --popGrowthModel unif exponential --stochasticGrowth t --
patchSize_file GilbertEtAl seq 1 100 2 seq 1e3 18e3 1e3
```

Some of the options have already been mentioned above. '--m LSS 3 0.3 0.4 0.3 1' sets a Linear Stepping Stone model, with 3 probability of events, where the probability of not migrating is the value at index 1 (zero based counting). It therefore sets a probability of migrating toward the left or right to 0.3 and the probability of not migrating to 0.4. By default, boundary conditions are dealt with by redistributing the probabilities of migrating toward the non-existence patch to the other probabilities (boundaries are reflective, not absorbing). '--fec 7' sets the fecundity for an individual having a fitness of one to 7. The keyword 'rep' used for option '--InitialpatchSize' works similar to the 'rep' function in R. Hence, 'rep 12 1e3' creates an array of the value 12 repeated 1000 times. Hence, '--InitialpatchSize A rep 12 1e3 rep 0 7e3' sets the initial patch size of the first 1000 patches to 12 and the initial patch size of the remaining 7000 patches to 0. '--popGrowthModel unif exponential' indicates that for all patches (hence the keyword 'unif' here) the growth model is an exponential model (bounded at the carrying capacity set by the option '--N'). '--stochasticGrowth t' indicates the growth must be stochastic ('t' is short for 'true'), that is the number of offspring in the next generation is sampled from a Poisson distribution with rate of the expected number of offspring as explained in chapter 3. The keyword 'seq' works like the homonym function in R. It creates an array starting from the first value, to the second value by the third value. For example, writing 'seq 0.1 0.9 0.2' is equivalent to writing '0.1 0.3 0.5 0.7 0.9'. '--patchSize_file GilbertEtAl seq 1 100 2 seq 1e3 18e3 1e3' asks for an output file indicating the patch size (number of individuals) of each patch. The file name is set to 'GilbertEtAl' (to which an automatic extension will be added) and output will be produced at every two generations from generation 1 to generation 100 and then every 1000 from 1000 to generation 18000.

For the simulation inspired from Booker and Keightley (2018), the input used is

```
SimBit --PN 1 --N A 1e3 --L T5 15e4 --T5_mu unif 2.5e-6 --T5_fit
cstHgamma 0.5 homo 0.111 2.22 --r rate unif 2.5e-6 --nbGens
20000
```

'`--T5_fit cstHgamma 0.5 homo  0.111 2.22`' indicates a Gamma distribution of homozygous (aka. double-mutant; hence the keyword 'homo') selection coefficient with parameters α = 0.111 and β = 2.22 and with a dominance coefficient of 0.5.

For the simulation inspired from O'neil et al. (2019), the input used was

```
SimBit --PN 3 --m island 0.000125 --N unif 1e3 --L t5 1e5 --
T5_fit multfitA rep 0.9995 25000 rep 1.0 75000 --r rate unif 0 -
-T5_mu unif 5e-07 --nbGenerations 2e4
```

'`--m island 0.000125`' indicates an island model (among the three patches defined with option '`--PN`') with a probability of migrating of 0.000125. '`--T5_fit multfitA rep 0.9995 25000 rep 1.0 75000`' sets the selection coefficients of the first 25000 loci to 1-0.9995=0.0005 and the selection coefficients of the remaining 75000 to 1-1=0 and indicate the use of the assumption of multiplicative fitness.

Finally, for the simulation inspired from the human genome and from ancestral human demography, the input used was

```
SimBit --PN 500 --N unif 100 --L T5 1e8 --T5_mu unif 1.25e-8 --r
rate unif 1e-8 --nbGenerations 1e3 --m LSS 1 0.1 0.8 0.1 --
T5_fit unif 1 0.99999
```

The above input does not use of the multiplicative fitness assumption. To make use of it, I replaced '`--T5_fit unif 1 0.99999`' with '`--T5_fit multfitUnif 0.99999`'.

SLiM and Nemo's input files tend to be much larger. As an example, for the simulation inspired from Gilbert et al. (2017), my input command for SimBit is 290 characters long (excluding the call of the executable) while the input file of Nemo is 55,698 characters long. I therefore only reported here, input examples for SimBit.
